# Distributed Genetic Effects on Human Brain Structure Emerge Across Multiple Spatial Scales

**DOI:** 10.64898/2026.08.17.745237

**Authors:** Emma J. Gleave, Luis M. García-Marín, Zuriel Ceja, Miguel E. Rentería, Tamoghna Chattopadhyay, Christian Gaser, Priya Rajagopalan, Paul M. Thompson

## Abstract

Genome-wide association studies (GWAS) have identified hundreds of common genetic variants associated with regional brain volumes, enabling the construction of polygenic scores (PGS) that summarize genetic predisposition for variation in specific neuroanatomical traits. To investigate how these genetic influences are exerted spatially throughout the brain, we computed PGS for ten brain volume phenotypes, including nine major subcortical structures and intracranial volume. Each locus was weighted by its estimated GWAS effect size on regional volume in the original GWAS. In an independent, non-overlapping sample of 2,830 UK Biobank participants, we performed whole-brain voxel-based morphometry (VBM) analyses of 3D volumetric brain MRI to reveal voxel-wise associations between each PGS and modulated gray matter volume (GMV). To probe genetic effects across multiple spatial scales, analyses were repeated across Gaussian smoothing kernels ranging from 2-mm to 12-mm full-width at half-maximum (FWHM). Several PGS demonstrated highly significant associations with GMV, including localized effects in the hippocampus, amygdala, thalamus, and basal ganglia, whereas the brainstem PGS showed more widespread associations throughout the brain. For most of the PGS, the fraction of voxels surviving the false discovery rate (FDR) correction increased with increasing FWHM. Peak voxel-wise significance was often strongest at intermediate smoothing levels. Hippocampal significance maps showed progressively larger regions of significant signal at higher smoothing levels, and subsampling showed that detectable signal remained present even with substantial reductions in sample size. These findings suggest that genetic influences on brain morphology are expressed across multiple spatial scales, with consequences that may help to guide the design of deep learning methods to discover genomic loci associated with brain structure and brain diseases.

## 1 Introduction

A central goal of imaging genetics [1] is to understand how genomic variation contributes to individual differences in the brain through genetic analyses of neuroimaging data, partly in an effort to understand how disease risk genes exert their effects. Large-scale genome-wide association studies (GWAS) have identified thousands of variants associated with neuroanatomical traits [2–4], enabling the construction of polygenic scores (PGS) that summarize genetic predisposition for variation in specific brain regions. Although PGS are typically derived for regional volumetric measures, less is known about how these influences are expressed spatially across the brain. For instance, *are the effects structure-specific, or do they extend to the rest of the brain?* Algorithms to map brain-wide effects of genetic polymorphisms have included voxel-wise GWAS [5], sparse PLS and CCA, parallel ICA, and reduced rank regression; deep learning methods to detect genetic effects on the brain include GWAS of latent space embeddings from autoencoders [6] and multimodal transformers [7]; the full range of current multivariate methods for imaging genomics are reviewed in [8]. While powerful, latent-space and embedding-based approaches can be difficult to interpret anatomically because associations are detected between complex image and genomic representations rather than with directly observable brain features. Voxel-based morphometry (VBM) may offer a more interpretable framework by relating genetic predictors to local gray matter volume (GMV) at voxel resolution. However, brain anatomy is a spatially organized system shaped by development, connectivity, and genetics rather than a collection of independent voxels. Data-driven and hierarchical clustering of genetic influences [9,10] suggest that genetic variants may affect distributed patterns of anatomical variation spanning multiple spatial scales, but most analyses were conducted in relatively small cohorts, limiting confidence in their reproducibility. Consistent with this, shared genetic influences can extend across anatomically distinct brain regions, perhaps even forming genetically organized networks that are not captured by local anatomical measures alone [10,11].

A key methodological consideration in VBM is the degree of spatial smoothing applied prior to statistical analysis. Smoothing is typically performed using a 3D Gaussian kernel defined by its full-width at half-maximum (FWHM) and is used to improve the signal-to-noise ratio, to reduce confounding effects of residual anatomical misregistration across individuals, and to better satisfy assumptions of voxel-wise statistical inference, such as the Gaussian random field theory used in the SPM analysis package [13,14]. Typical smoothing kernel sizes range from 2- to 12-mm FWHM depending on the anatomical structures under investigation and the hypothesized spatial extent of the effects (based on the matched filter theorem) [7,12]. The default smoothing kernel in Statistical Parametric Mapping (SPM) is 8-mm FWHM to provide a compromise between anatomical specificity and statistical sensitivity [14,15]. Previous VBM sensitivity analyses show that larger smoothing kernels tend to produce more spatially extensive findings, with gray matter effects becoming increasingly widespread as kernel size increases [15]. In any voxel-based brain mapping method, the structures of interest vary in size and spatial organization. Smaller kernels may preserve local anatomical detail but larger kernels may improve sensitivity at the cost of spatial specificity [15,16-18]. If genetic effects are coherent across broader anatomical regions rather than isolated voxels, different smoothing scales may reveal different components of the underlying signal, making kernel choice informative about the spatial scale of genetic effects, and not merely a preprocessing decision.

Here, we examined how smoothing kernel size affects sensitivity to voxel-wise imaging-genetics associations and sought parameter settings that may help to improve future genomic discovery. Using polygenic scores for major subcortical structures, we performed whole-brain VBM analyses across Gaussian kernels from 2-mm to 12-mm FWHM and compared significance distributions, voxel-wise maps, and the fraction of voxels surviving false discovery rate (FDR) correction. We asked whether genetic effects on brain morphology would vary with scale and whether smoothing trajectories can reveal the spatial coherence of genetic influences on the brain.

## 2 Methods

### Dataset and MRI Acquisition

The primary cohort for our PGS-VBM analyses was drawn from UK Biobank (UKB), a longitudinal prospective population study of over 500,000 middle-aged to older adults, with extensive health, demographic, imaging, and genetic data collected across the United Kingdom [19,20]. Available data includes T1-weighted (T1w) brain MRI, genome-wide genetic data, and pre-computed genetic principal components (PCs) summarizing ancestry-related population structure. These PCs are commonly incorporated as covariates in genomic analyses [21,22]. In our study, the first four PCs were included in all statistical models. To ensure sample independence, we excluded participants included in the discovery GWAS of Garcia-Marin et al. (2024) [2], avoiding circularity between SNP discovery and downstream imaging analyses. Inclusion required the availability of a baseline T1w MRI, genome-wide genetic data, and demographics (age and sex). Participants with missing or quality-control–failed data (e.g., motion artifacts) were excluded. The final sample comprised 2,830 individuals (age: 68.8 ± 8.1 years; 51.0% female) scanned across four UKB imaging sites.

### Statistical Framework for PGS Derivation

Ten regional brain volume phenotypes were evaluated: nine subcortical regions of interest (ROIs)—the amygdala, caudate nucleus, hippocampus, nucleus accumbens, globus pallidus, putamen, thalamus, ventral diencephalon, and brain stem— and a global measure of intracranial volume (ICV). For each ROI, a PGS was calculated by weighting genetic loci by their effect size estimates. Specifically, for individual (*i*) and ROI (*r*), the score was calculated as:

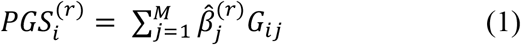

where *j* indexes single-nucleotide polymorphisms (SNPs), *M* is the total number of SNPs contributing to the score, *G*_*ij*_ represents the number of effect alleles (0, 1, or 2) carried by individual *i* at locus *j*, and 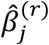 is the estimated effect of SNP *j* on ROI *r*. Polygenic scores were generated using SBayesR [23] within the Genome-wide Complex Trait Bayesian (GCTB v2.5.4) framework, with the UK Biobank 7-million imputed SNP linkage disequilibrium reference panel [24]. SBayesR jointly estimates SNP effects while accounting for linkage disequilibrium and the highly polygenic architecture of brain-volume traits, producing shrinkage-adjusted effect estimates that improve predictive performance. The resulting PGS is an aggregate measure of genetic influences on variation on a specific brain structure.

### Image Processing and FWHM Overview

To investigate the anatomical distribution of these polygenic effects, voxel-based morphometry (VBM) was used to quantify modulated GMV at each voxel throughout the brain [25]. T1w scans and gray matter segmentations were processed using the ENIGMA Computational Anatomy Toolbox (CAT12) (**Fig. 1**; https://neuro-jena.github.io/enigma-cat12/) [26], implemented within the Statistical Parametric Mapping (SPM) MATLAB framework [27]. This pipeline performs voxel-wise tissue segmentation via a series of standardized processing steps including denoising, bias-field correction, tissue classification, and spatial registration. CAT12 produces voxel-wise gray matter and white matter tissue masks, along with regional and global volumetric estimates including gray matter, white matter, cerebrospinal fluid (CSF), and total intracranial volume (TIV). During spatial normalization, gray matter tissue maps were modulated using the Jacobian determinants of the deformation fields, thereby preserving local volumetric information prior to statistical analysis. Following segmentation and registration, gray matter maps were smoothed using 3D Gaussian kernels with FWHM values of 2-mm, 4-mm, 6-mm, 8-mm, 10-mm, and 12-mm.

**Fig. 1.**
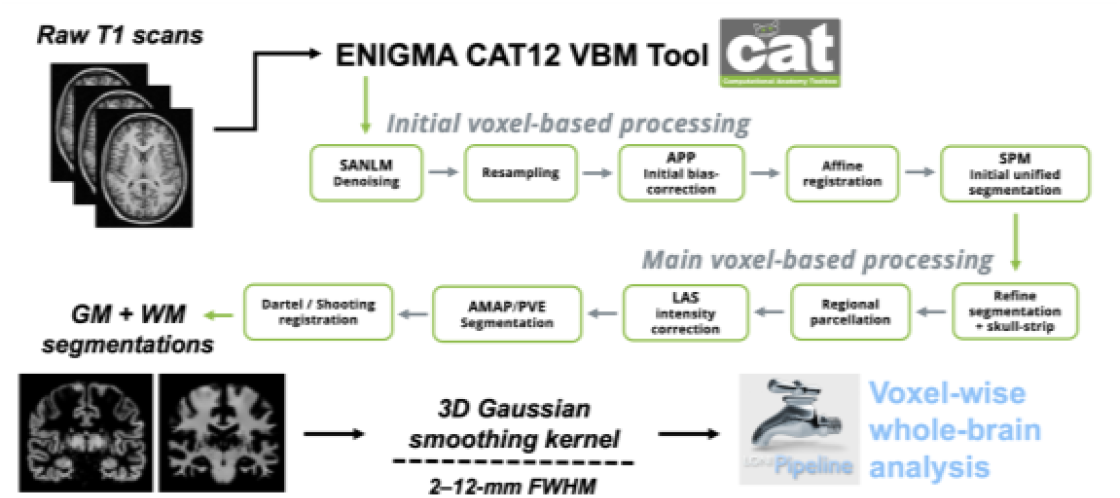
VBM Pipeline and Analysis Workflow. The ENIGMA CAT12 VBM pipeline [26] processes raw 3D T1-weighted MRI scans in two stages. The first applies spatial adaptive non-local means (SANLM) denoising, bias correction, segmentation, and registration. The second performs skull stripping, regional parcellation, intensity normalization, and refinement of tissue classification and volume estimates for voxel-wise analysis. LAS (Local Adaptive Segmentation) improves local tissue intensity normalization, AMAP (Adaptive Maximum A Posteriori) segmentation classifies tissue without population priors, and PVE (Partial Volume Estimation) accounts for voxels containing mixtures of tissue types, improving accuracy at tissue boundaries.

### Statistical Analysis and FWHM Optimization

The relationship between voxel-wise GMV and each ROI-specific PGS was evaluated using a linear mixed-effects model (**Fig. 1**; LONI Pipeline [http://pipeline.loni.usc.edu] v.7.0.3; R version 4.5.0) [28]:

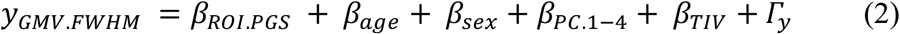

where age, sex, the first four UKB ancestry PCs, and TIV were included as fixed effects, while scanner site was modeled as a random effect *y* to account for site-specific variability across UKB acquisition centers. Separate voxel-wise regressions were performed for each ROI PGS at each smoothing size. Multiple-comparison correction was performed using the standard false discovery rate (FDR) procedure at (*q*=0.05) [29]. To limit multiple testing across smoothing scales, the 2-mm FWHM analysis, which offers the finest spatial resolution, served as the primary analysis, while the remaining smoothing levels were considered to be exploratory. These additional scales provide different views of the underlying anatomical signal rather than truly independent confirmatory tests. Alternatively, FDR could be controlled across all voxel-by-scale comparisons with a scale-space multiple-testing procedure.

To understand how the detectability of genetic effects depends on spatial scale, we quantified three complementary properties of the voxel-wise association maps: (i) the global departure from the null distribution using quantile-quantile (QQ) plots, (ii) the voxel-wise significance threshold after false discovery rate (FDR) correction, and (iii) the spatial extent of significant effects, measured as both the number of voxels surviving FDR correction (FDRvox) and the fraction of voxels surviving correction within the common template-space analysis mask (FracFDR), which comprised 382,592 voxels and was identical across all subjects and analyses. These measures summarize the strength, extent, and statistical significance of the genetic signal across smoothing scales. To summarize the influence of smoothing on detection sensitivity, FracFDR values obtained from the full sample were plotted as a function of FWHM for all PGS analyses (**Fig. 3**). Voxel-wise significance maps were generated from the FDR-corrected *p*-values of each ROI PGS analysis. Thresholded *p*-values were transformed to a −log_10_(p) scale and displayed with a common colorbar to facilitate comparison across smoothing levels; representative results for hippocampus shown in **Fig. 4**. To assess the effect of sample size on detection sensitivity, we show hippocampus PGS analyses at both 2-mm and 12-mm FWHM (n=2,830) subsampled to 75% (n=2,122), 50% (n=1,415), and 25% (n=707), with 35 independent replicates generated at each sampling level. FracFDR was plotted for each replicate across sample sizes with the median and bootstrap 95% confidence interval of the median (10,000 bootstrap resamples) shown **(Fig. 5)**.

**Fig. 2.**
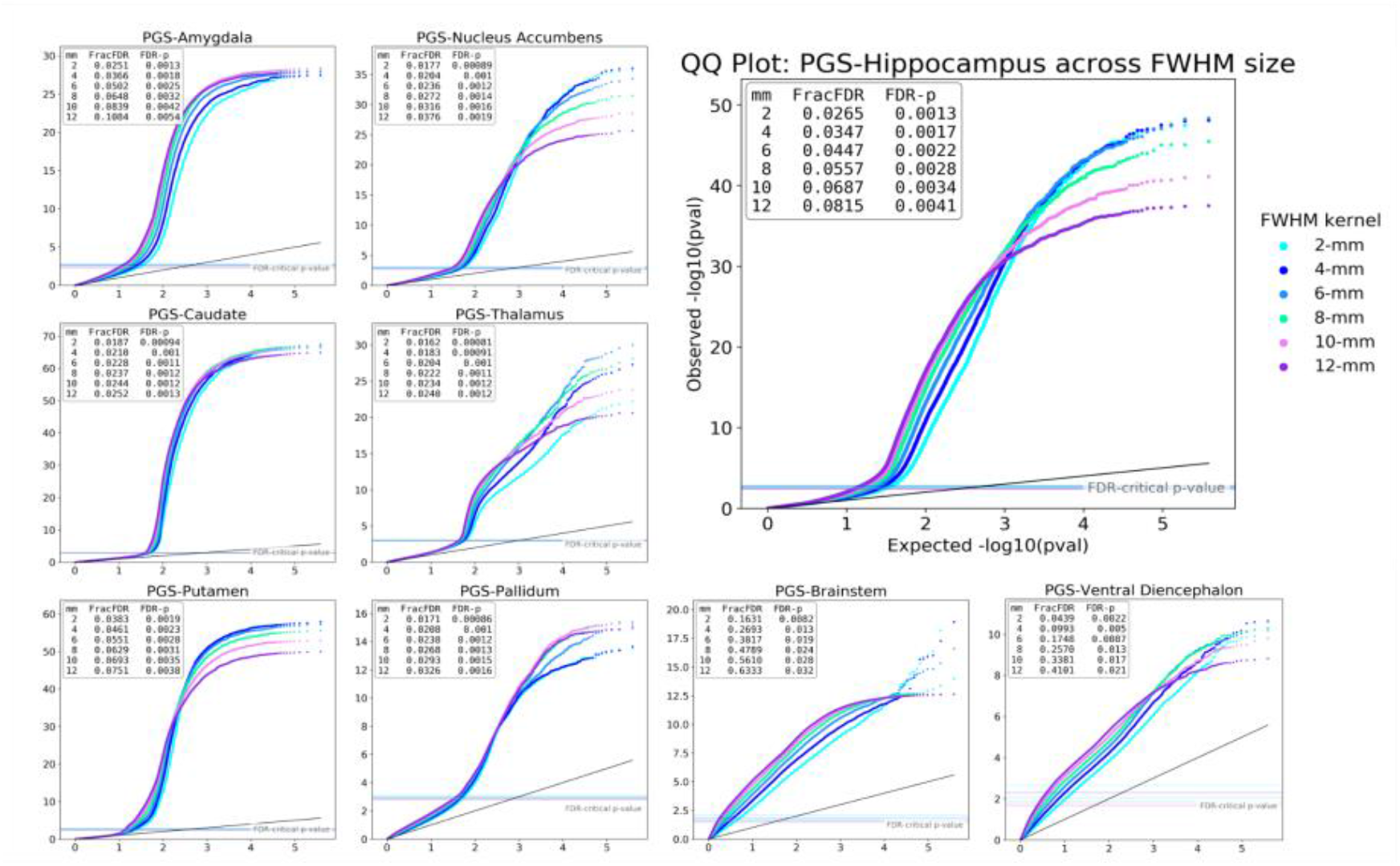
Quantile-Quantile (QQ) plots: significance values for the brain-wide statistical effect of each subcortical PGS by FWHM kernel size. QQ plots of raw p-values from whole-brain voxel-wise linear mixed regression models for the association between polygenic score for nine brain ROIs (hippocampus, amygdala, nucleus accumbens, caudate, thalamus, putamen, pallidum,n brainstem, & ventral diencephalon) and GMV across 2-mm to 12-mm kernel sizes. FDR-correction (*q*=0.05) *p*-value thresholds are plotted on the *x*-axis for each ROI FWHM analysis set, and summarized per plot box as “FDR-p” alongside FracFDR.

**Fig. 3.**
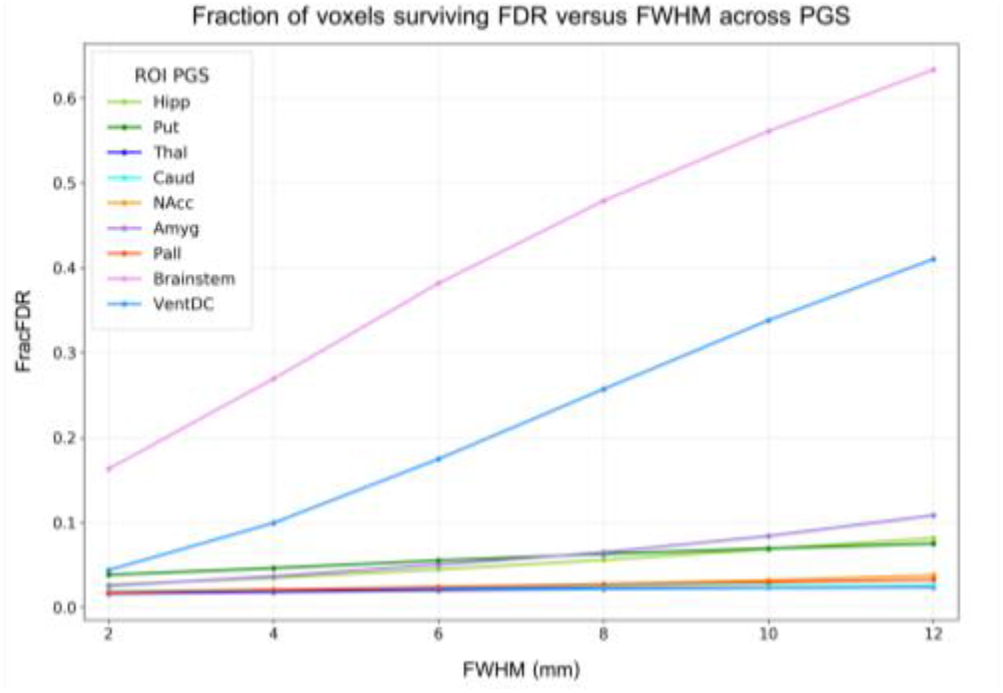
Fraction of voxels surviving FDR from each subcortical ROI PGS analysis at full size (n=2,830) across all FWHM kernel sizes. The statistical maps of brain-wide associations with nine ROI PGS plotted include: the hippocampus, putamen, thalamus, caudate, nucleus accumbens, amygdala, pallidum, brainstem, and ventral diencephalon, and from analyses across all 3D Gaussian FWHM smoothing kernel sizes, 2-mm to 12-mm, in 2-mm increments.

**Fig. 4.**
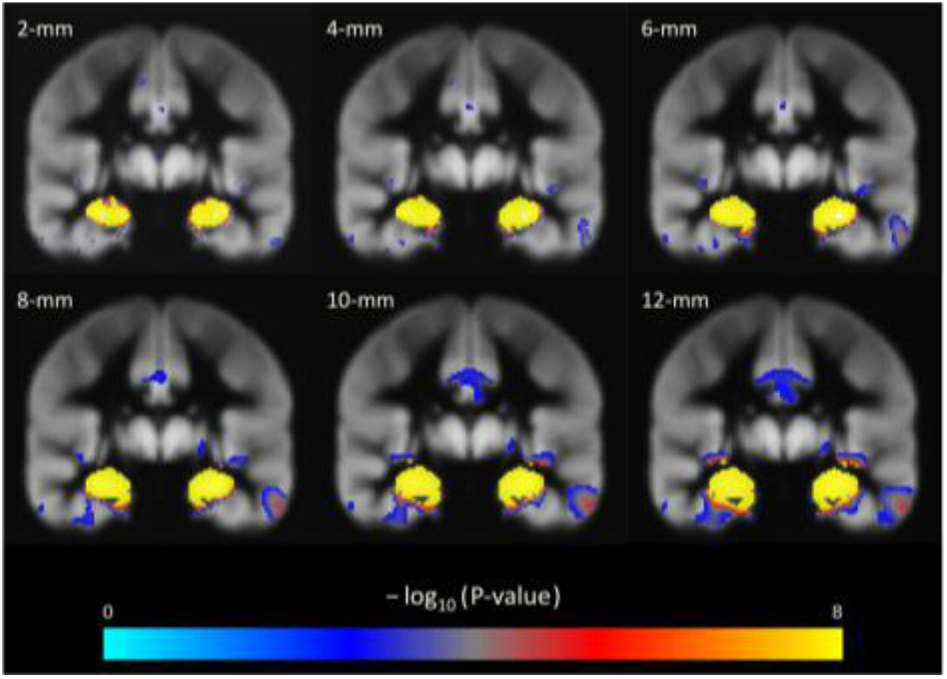
Voxel-wise significance maps for whole-brain GMV associations with the PGS for hippocampal volume, across kernel sizes. FDR-corrected (*q*=0.05) significance maps show full-sample (n=2,830) results at FWHM smoothing kernel sizes: 2-mm (*top left*; standard-FDR critical *p*-value=0.0013), 4-mm (*top center*; standard-FDR critical *p*-value=0.0017), 6-mm (*top right*; standard-FDR critical *p*-value=0.0022), 8-mm (*bottom left*; standard-FDR critical *p*-value=0.0028), 10-mm (*bottom center*; standard-FDR critical *p*-value=0.0034), and 12-mm (*bottom right*; standard-FDR critical *p*-value=0.0041). Maps were log transformed and thresholded. Coronal slices are shown at the median peak Y coordinate across FWHM kernels (Y=-18 mm).

**Fig. 5.**
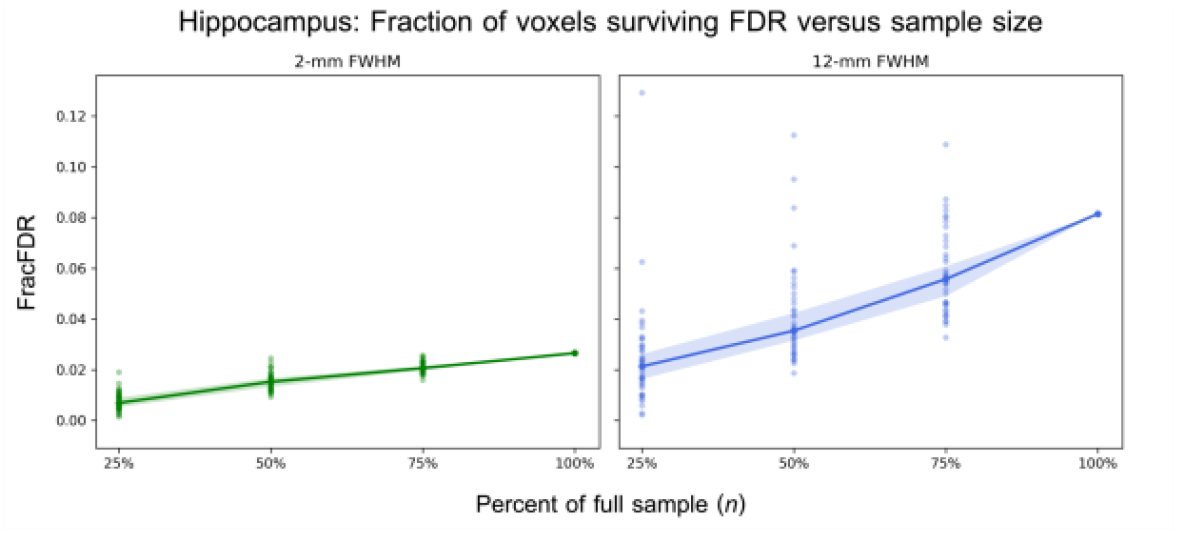
Fraction of voxels surviving FDR in the analyses of the PGS for hippocampal volume at 2-mm vs 12-mm across sample sizes. Plots for both 2-mm and 12-mm FWHM display FracFDR versus sample size from subsampling of the PGS hippocampus analyses at full size (n=2,830), then 75% (n=2,122), 50% (n=1,415), and 25% (n=707) sample size replicated 35 times each per FWHM analysis, with the median FracFDR line and shaded bootstrap 95% confidence interval of the median shown.

## 3 Results

QQ plots across all smoothing levels (**Fig. 2**) showed that in general, top *p*-values weakened slightly at larger FWHM values, although they remained highly significant across all smoothing levels. The strongest deviation from the theoretical null occurred around 4–6-mm FWHM for the hippocampus, while FracFDR still continued to increase from approximately 6.8% to 8.2% at the largest kernels. Similar patterns were observed for the nucleus accumbens, putamen, and thalamus, where the upper tail of the QQ distribution was strongly elevated relative to the theoretical null while FracFDR increased with smoothing. Across all subcortical ROI analyses **(Fig 3)**, FracFDR generally increased with increasing FWHM and was particularly pronounced in the brainstem and ventral diencephalon, increasing several-fold between 2-mm and 12-mm FWHM. Hippocampus, amygdala, putamen, and thalamus showed more moderate but still progressive increases across smoothing levels.

Coronal slices of the hippocampal PGS significance maps (**Fig. 4**) show a steady growth of FDR-significant signal with increasing FWHM. Significant voxels are already visible at 2-mm FWHM but extend over a broader area at higher smoothing levels, especially between 8-mm and 12-mm. While the size of the signal changes markedly, its primary hippocampal location is largely unchanged across kernels. FracFDR rose steadily with sample size for both kernels (**Fig. 5**), peaking in the full cohort (n=2,830). The 12-mm FWHM analyses retain a larger fraction of FDR-significant voxels than the 2-mm analyses at every sample size tested. Significant signal was still detectable in the smallest subsample (n=707), though fewer voxels survived FDR correction than in larger samples. Bootstrap confidence intervals around the median FracFDR demonstrated increasing precision in the estimated detection sensitivity with increasing sample size.

## 4 Discussion

Across QQ plots, significance maps, FracFDR trajectories, and subsampling analyses, the results show that a substantial part of the PGS-related anatomical signal is distributed across space rather than confined to isolated voxels. As FWHM increased, FracFDR generally rose across most structures, while hippocampal maps showed a steady growth in FDR-significant voxels. QQ plots revealed that the strongest voxel-wise effects were often retained at intermediate smoothing levels. Overall, these findings imply that genetic effects on brain structure are scale-dependent, and different features emerge at different smoothing levels. Rather than reflecting isolated voxel-wise associations, the signal tends to lie in broad anatomical patterns that become easier to detect as fine-scale variation is suppressed. The steady rise in FracFDR and expansion of significant hippocampal signals at large kernels support the view that genetic influences on brain morphology may be organized as coherent anatomical modes.

These findings may help to design new multivariate, multiscale models for imaging genetics. The patterns seen across scales suggest that genes may shape the brain through a set of coherent anatomical patterns. We can estimate these by building a genetic correlation matrix between voxels or regions and extracting its leading eigenmodes. The same matrix may be viewed as a graph, whose Laplacian eigenvectors define smooth anatomical patterns. This would replace millions of voxel-wise tests with a set of genetically informed modes. Future models could learn these modes and their genetic drivers; in the same vein, transformers could serve as a nonlinear form of kernel CCA that links genes and brain structure through shared information. Such representations may explain a greater proportion of brain variation than mass-univariate methods, revealing local effects and broader genetic covariance. In future work, iteratively removing dominant modes may help to discover a hierarchy of independent genetic influences.

### Limitations

Analyses were based on modulated, spatially normalized gray matter images and currently only reflect gray matter volume variation. Several structures examined, including the caudate and thalamus, do contain some white matter, and structural differences in these regions may be only partly captured by gray matter VBM, reducing estimated effect sizes. Future analyses with deformation-based morphometry (DBM) and CAT12-derived Jacobian determinant maps will provide a more complete measure of these effects.

## 5 Conclusion

In an independent sample of 2,830 UK Biobank participants, we mapped the voxel-wise anatomical coherence of the statistical effect of polygenic scores derived for major subcortical brain structures. Smoothing kernel size influenced statistical maps, QQ plots of the statistical effects, and the fraction of voxels surviving FDR correction. The strongest voxel-wise effects were often observed at intermediate smoothing levels, but larger FWHM kernels generally increased the spatial extent of significant signals across most structures. These analyses suggest that a substantial component of the genetic signal underlying brain structural variation is spatially distributed and expressed across multiple anatomical scales. Rather than treating smoothing as a preprocessing choice, a scale space search may help to discover the underlying genomic signal at currently available sample sizes. We will use these findings to guide the design of multiscale transformer models with the goal of explaining greater variance in brain maps.

## Acknowledgements

Supported by NIA grants R01AG058854 (ENIGMA World Aging Center), U01AG068057 (AI4AD), and an Alzheimer’s Association grant AACSFD-22-974694 (to P.R.).

